# Education differentiates cognitive performance and resting state fMRI connectivity in healthy aging

**DOI:** 10.1101/2023.02.22.529483

**Authors:** Sonia Montemurro, Nicola Filippini, Giulio Ferrazzi, Dante Mantini, Giorgio Arcara, Marco Marino

**Author notes:** **Corresponding author:** Marco Marino, Movement Control & Neuroplasticity Research Group, Tervuursevest 101, box 1501, 3001 Leuven, Belgium. these authors share first autorship (Sonia Montemurro, Nicola Filippini). these authors share last autorship (Giorgio Arcara, Marco Marino). **DECLARATION OF INTERESTS:** None.

## Abstract

**Objectives:** In healthy aging, the way people differently cope with cognitive and neural decline is influenced by the exposure to cognitively enriching life-experiences. Education is one of them, so that in general the higher the education the better the expected cognitive performance in aging. At the neural level, it is not clear yet whether education can differentiate resting state functional connectivity profiles and their cognitive underpinnings. Thus, with this study, we aimed to investigate whether the variable education allowed for a finer description of age-related differences in cognition and resting state FC.

**Methods:** We analyzed in 197 healthy individuals (137 young adults aged 20-35, and 60 older adults aged 55-80 from the publicly available LEMON database), a pool of cognitive and neural variables, derived from magnetic resonance imaging, in relation to education. Firstly, we assessed age-related differences, by comparing young and older adults. Then, we investigated the possible role of education in outlining such differences, by splitting the group of older adults based on their education.

**Results:** In terms of cognitive performance, older adults with higher education and young adults were comparable in language and executive functions. Interestingly, they had a wider vocabulary compared to young adults and older adults with lower education. Concerning functional connectivity, the results showed significant age- and education-related differences within three networks: the Visual-Medial, the Dorsal Attentional and the Default Mode network (DMN). For the DMN, we also found a relationship with memory performance, which strengthen the evidence that this network has a specific role in linking cognitive maintenance and FC at rest in healthy aging.

**Discussion:** Our study revealed that education contributes to differentiate cognitive and neural profiles in healthy older adults. Also, the DMN could be a key network in this context, as it may reflect some compensatory mechanisms relative to memory capacities in older adults with higher education.

## 1. Introduction

Age-related changes on cognitive function may influence the quality of life (Harada et al., 2013; Murman, 2015). For example, memory performance worsens with age and well-maintained memory capacities could be considered as a “tract” of a preserved cognitive function in healthy aging (Harada et al., 2013; Nyberg and Pudas, 2019). Conversely, some cognitive abilities, such as the capacity to retain words in one’s own vocabulary (Verhaeghen, 2003), are resistant to aging and they may even improve across life (Murman, 2015). In this context, the concept of cognitive reserve (CR, Lojo-Seoane et al., 2018) may help to explain differences in the way people get older and cope with different cognitive demands, which depend on the cognitive “resources” accrued during life (Barulli and Stern, 2013). Interestingly, although many other proxies like occupational attainment or IQ may determine CR (Lojo-Seoane et al., 2018; Montemurro, 2022), education is one of the mostly used. Indeed, it influences late-life cognitive function by contributing to individual differences in cognitive skills (Lovden et al., 2020), where the higher the education, the better the expected cognitive performance in aging (Elkins et al., 2006; Montemurro et al., 2019; Montemurro, 2019). This is also in line with recent research from our group reporting that education represents a protective factor against age-related cognitive decline (Mondini et al., 2022). In addition, education is one of the most accessible and suitable CR proxies (e.g., easy to collect) for aging populations (Chapko et al., 2018; Montemurro et al., 2021), besides being one of the most commonly used in clinical and experimental contexts (Meng and D’Arcy, 2012; Anaturk et al., 2021).

Whilst education has been so far mainly used to explain cognitive decline in aging as a confound (i.e., nuisance) or a predictor (i.e., proxy), a full comprehension of its role in outlining brain functioning and its relationship with cognitive function still remains challenging. Brain aging has been investigated through magnetic resonance imaging (MRI) in terms of structural changes (e.g., atrphy of cortical grey matter or increased cerebrospinal fluid volume, (Reuter-Lorenz, 2002; Raz et al., 2005; Fjell et al., 2009; Park and Reuter-Lorenz, 2009; Salthouse, 2011; Orellana et al., 2016; Oschwald et al., 2019), and functional changes (Vidal-Pineiro et al., 2014; Yoshimura et al., 2020), showing that some brain networks, like the Default Mode Network (DMN), are affected by aging. Functional connectivity, measured with functional MRI (fMRI), allows to investigate the relationship between fluctuations among different brain areas (Sporns, 2013). This technique has been largely used to study brain networks in aging (Vidal-Pineiro et al., 2014; Yoshimura et al., 2020) and their behavioral correlates (Farras-Permanyer et al., 2019; Varangis et al., 2019). For example, it has been shown that brain network integrity provides a helpful outlook on the preservation of cognitive functioning (Geerligs et al., 2015; Jockwitz and Caspers, 2021), and that such integrity is mainly investigated by resting state networks (RSNs), whose topology recalls the ones of brain networks emerging during active tasks (Smith et al., 2009). Changes in RSNs have been linked to cognitive decline in aging, but a high heterogeneity has limited the understanding of the impact of aging on brain function.

In this context, CR has gained relevance and has been integrated in neuroimaging research (Bastin et al., 2012; Menardi et al., 2018). CR should be ideally helpful for a better comprehension of individual differences in aging, by using proxies that allow to estimate the individuals’ potential resilience. However, this field of research is still presenting challenges. Several theories have been already proposed to explain functional changes in aging, i.e., the Hemispheric Asymmetry Reduction in Older Adults model (HAROLD, Cabeza, 2002), the Posterior-Anterior Shift in Aging theory (PASA, Davis et al., 2008) the Compensation-Related Utilization of Neural Circuits Hypothesis CRUNCH, Reuter-Lorenz, 2008), and the Scaffolding Theory of Aging and Cognition (STAC, Park and Reuter-Lorenz, 2009). Among them, the STAC, lately revised in the STAC-r (Reuter-Lorenz and Park, 2014), introduced education (and other CR proxies) as a crucial variable to interpret the mechanisms of aging (Nyberg et al., 2012). In STAC-r, a high level of education is conceived as being associated with an enhanced neurocognitive scaffolding, so that, despite neural deterioration, cognitive function should be maintained in older adults (Reuter-Lorenz and Park, 2014). Nevertheless, the role of education in determining cognitive and brain characteristics in healthy aging populations has also displayed some diverging findings (Nyberg et al., 2012), and although education has been much discussed such a factor that may better explain inter-individual differences in aging, it is still unclear how neurocognitive profiles could be characterized based on differences in such an important variable.

In the present study, starting from the premise that, in healthy aging, cognitive performance is better maintained with a higher education, we tested whether education could differentiate healthy older adults in terms of not only cognitive performance but also fMRI connectivity, especially for RSNs involved in high-level cognitive functions. To this end, we first compared older adults to a more preserved population, i.e., younger adults, to identify age-related differences. Then, by splitting the older adult group in two subgroups based on education (higher and lower), we specified the effect of education in healthy aging. Accordingly, we expected that cognitive and neural variables would show age-related differences and more importantly, at the basis of this study, that education would provide definite information about age-related differences in resting state fMRI connectivity and its relationship with cognition.

## 2. Material and methods

### 2.1. Participants and materials

All participants included in this study were taken from the publicly available database “Leipzig Study for Mind-Body-Emotion Interactions” (LEMON) (Babayan et al., 2019). Data collection was performed in accordance with the Declaration of Helsinki, approved by the local ethics committee and all participants provided written informed consent prior to data acquisition for the study (including agreement to their data being shared anonymously). Following the application of exclusion criteria, which are listed in detail in Babayan et al. 2019 (Babayan et al., 2019), the total sample included 227 participants. Due to missing data of specific cognitive variables of interest, 30 participants were excluded from the analysis, thus the total number of participants was n=197. The subjects were separated into two groups, based on their age, resulting in one group of younger adults (young, Y, n = 137, range 20-35 years) and in one of older adults (old, O, n = 60, range 55-80 years). The old group was further divided into two subgroups based on their education: higher education (old-high, OH, n = 30), i.e., *lyceum/gymnasium* (12 years), and lower education (old-low, OL, n = 30), i.e., *technical high school/realschule* (10 years). Notably, all younger adults had higher education, as the old-high group. Five cognitive tests addressing memory, language, and vocabulary were used. The behavioral tasks included: *a) Short-* and *b) Long-term Memory tests* (*California Verbal Learning Task, CVLT, Niemann, 2008)*, used to assess verbal learning and memory capacity, and to provide information about different learning strategies by testing Immediate Memory Recall and Delayed Memory Recall; *c) Vocabulary test* (*Wortschatztest, WST, Schmidt, 1992)*, used to measure verbal intelligence, and assess language comprehension; *d) Phonemic* and *e) Semantic Fluency tests* (*Regensburger Wortflüssigkeitstest, RWT, Aschenbrenner, 2000)*, used to assess verbal fluency. Notably, higher scores correspond to a better cognitive performance in all cognitive tests. A full description about the cognitive assessment can be found in Babayan et al. (Babayan et al., 2019). Demographic variables, such as sex and Body Mass Index (BMI) were also assessed.

### 2.2. MRI data acquisition

MRI scans were performed on a Siemens 3 Tesla (3T) MAGNETOM Verio MR scanner (Siemens Healthcare GmbH, Erlangen, Germany) equipped with a 32-channel receiver head coil. Briefly, the data used in this study included anatomical and resting state fMRI (rs-fMRI) scans. The anatomical scans were acquired using a 3D T1-weighted (T1w) Magnetization-Prepared 2 Rapid Acquisition Gradient Echoes sequence (MP2RAGE) (Marques et al., 2010), with a MP2RAGE block time = 5000ms, a repetition time (TR) = 6.9ms, an echo time (TE) = 2.92ms, inversion time (TI1) = 700ms, TI2 = 2500ms, flip angle (FA1) = 4°, FA2 = 5°, voxel dimension = 1 mm isotropic, acquisition time = 8 min 22s. The rs-fMRI scans were acquired using a T2*-weighted echo planar imaging (EPI) sequence with a multiband factor MB = 4, TR = 1400ms, TE = 30ms, FA = 69°, voxel dimension = 2.3mm isotropic, number of volumes (N) = 657 volumes, acquisition time = 15min 30s. The two images acquired at TI1 and TI2 are fused into a unique anatomical image with enhanced T1 contrast and reduced bias field.

During the rs-fMRI run, participants were instructed to remain awake and lie down with their eyes open while looking at a low-contrast fixation cross. Spin echo EPI with reversed phase encoding were also acquired and used for rs-fMRI distortion correction. These scans were acquired with TR = 2200 ms, TE = 50 ms, FA = 90°, voxel dimension = 2.3 mm isotropic, phase encoding = anterior to posterior (AP) and posterior to anterior (PA), acquisition time = 29s each. Further details about the MRI protocol can be found in Babayan et al. (Babayan et al., 2019). To quantify eventual head motion artifacts, we calculated the framewise displacement (FD, Power et al., 2012) computed as the sum of the absolute values of the derivatives of the translational and rotational realignment estimates at every timepoint, for which we reported values below 0.5 for all groups (FD_HC_ = 0.18 ± 0.06, FD_OH_ = 0.22 ± 0.14, FD_OL_ = 0.23 ± 0.09).

### 2.3. Data processing

#### 2.3.1. MRI data analysis

##### Anatomical scans

Voxel-Based-Morphometry (VBM) was used to study GM differences between young and older adult groups. Preprocessing was carried out using FSL (www.fmrib.ox.ac.uk/fsl) and included: brain extraction and brain tissues segmentation (using FAST), which allows extracting measures of the total gray matter (GM), the white matter (WM) and the cerebrospinal fluid (CSF). Brain structural imaging features, including total brain volume (TBV) and segmented GM, WM, and CSF, were computed for each participant. Whole-brain analysis was carried out with FSL-VBM (Douaud et al., 2007), using default settings. In brief, brain extraction and tissue-type segmentation were performed and resulting GM partial volume images were aligned to the standard space using FMRIB’s Linear Image Registration Tool (FLIRT) and then nonlinear (FNIRT) registration tools. The resulting images were averaged, modulated, and smoothed with an isotropic Gaussian kernel of 3 mm to create a study-specific template. Finally, voxel-wise GLM was applied using permutation nonparametric testing (5,000 permutations), correcting for multiple comparisons across space.

##### rs-fMRI scans

Motion correction was performed using MCFLIRT, included in the fMRI Expert Analysis Tool (FEAT) toolbox (Woolrich et al., 2001). Then, distortion correction was performed using the fMRI datasets acquired with AP and PA phase-encoding directions. These were used to estimate a susceptibility induced off-resonance distortion field (Smith et al., 2004). The computed field was used to correct the, previously motion-corrected, time-averaged fMRI dataset. Following this, rigid registration of the motion and distortion corrected time-averaged fMRI data onto the individual structural T1w data was performed. After linear registration to the T1w image was performed to register the rs-fMRI data from the individual space to the Montreal Neurological Institute (MNI) standard space, a non-linear transformation was computed by processing the anatomical T1w image alone. In particular, the T1w image was registered to the MNI space in two sequential steps. First, an affine transformation (12 degrees of freedom) was computed. Then, a further non-rigid transformation was employed to reach fine-grained alignment. Note that, in this context, the affine transformation was employed as seed for the minimization of the non-linear registration step. After all these steps were performed, the off-resonance distortion field, the linear transformation mapping the fMRI data onto the T1w image, and the non-linear transformation mapping the T1w data onto the MNI space, were combined into an unique warping field, which was applied at once to the motion and distortion corrected fMRI time-series. After alignment was achieved, the fMRI data on the MNI space was finally spatially smoothed using a Gaussian kernel of full-width at half-maximum (FWHM) of 5 mm, and a high-pass temporal filtering equivalent to 100 s applied. fMRI connectivity analysis at rest was carried out using the Multivariate Exploratory Linear Optimized Decomposition into Independent Components (MELODIC) (Beckmann and Smith, 2004). Preprocessed functional data containing N = 657 volumes for each subject were temporally concatenated across subjects to create a single 4D dataset. The between-subject analysis of the rs-fMRI data was carried out using the “dual regression” approach (Filippini et al., 2009) that allows for voxel-wise comparisons of resting state functional connectivity. In this analysis, the dataset was decomposed into 25 components, in which the model order was estimated using the Laplace approximation to the Bayesian evidence for a probabilistic principal component model. The resulting group-ICA (Independent Component Analysis) components were visually inspected and then labelled as resting state networks (RSNs) based on the topology of their thresholded spatial maps. Thirty subjects for each of the three groups were considered to create the group-components. The mean connectivity values associated with each subject RSN maps were extracted and used as a measure of connectivity strength, referring to the spontaneous and synchronous activity within a given network for a specific subject. Statistical analysis, described in more detail in the following section (2.3.2), consisted of an Analysis of Covariance (ANCOVA), including TBV, BMI and sex as nuisances, which was run to test for differences in neural variables between the different groups.

#### 2.3.2. Statistical Analyses

Statistical analyses on continuous variables (i.e., socio-demographic, cognitive scores and brain measures) and dichotomic variables (i.e., sex) were carried out using SPSS software (SPSS, Inc.). A t-test was used to compare those variables between younger and older participants, whereas the Analysis of Variance (ANOVA) was used to compare socio-demographic variables and structural brain measures between the young, old-high, and old-low groups. The ANCOVA was then used to compare cognitive scores and RSN measures among all groups. The ANCOVA analysis included TBV, BMI and sex as nuisances (Table 1). Both ANOVA and ANCOVA post-hoc analyses were Bonferroni-corrected. Pearson’s correlation coefficients between cognitive scores and connectivity strength of RSNs that revealed significant group differences were calculated through a partial correlation analysis including TBV, BMI and sex as nuisances. The Fisher’s-r-to-Z transformation of the Pearson’s correlation was used to compare the partial correlation coefficients.

**Table 1.** Descriptive analyses and Analysis of Variance (ANOVA) of socio-demographic variables, and brain measures for all the groups, i.e. younger (Y), older (O), older with higher level of education (OH), and older with lower level of education (OL). The table shows for each variable mean and standard deviation or number of subjects. Columns show the values of the descriptive statistics for each group separately, i.e., young, old, old-high, and old-low, and for each comparison between groups, i.e. young vs old (third column), young vs old-high (seventh column), young vs old-low (eighth column), and old-high vs old-low (ninth column) contrasts. The Bonferroni-corrected p-values associated with the one-way ANOVA (sixth column) are reported for each contrast. Asterisks indicate significant results.

|  | Y<br>(N=137) | O<br>(N=60) | Y vs O<br>*p-val* | OH<br>(N=30) | OL<br>(N=30) | *p*-ANOVA | Post-Hoc comparisons<br>(Bonferroni-corrected) |  |  |
| --- | --- | --- | --- | --- | --- | --- | --- | --- | --- |
|  |  |  |  |  |  |  | Y vs OH | Y vs OL | OH vs OL |
| Socio-Demographic variables |  |  |  |  |  |  |  |  |  |
| Age | 22.83±3.42 | 64.50±4.84 | <0.001* | 64.16±5.88 | 64.83±3.59 | <0.001* | <0.001* | <0.001* | 1 |
| Sex | 93 M<br>44 F | 30 M<br>30 F | 0.02* | 18 M<br>12 F | 12 M<br>18 F | 0.006 | - | - | - |
| BMI | 23.01±2.92 | 26.31±4.12 | <0.001* | 26.51±4.08 | 26.10±4.22 | <0.001* | <0.001* | <0.001* | 1 |
| Brain Measures |  |  |  |  |  |  |  |  |  |
| Total Brain Volume | 1.57±0.01 | 1.49±0.02 | <0.01* | 1.55±0.03 | 1.44±0.03 | <0.001* | 1 | <0.001* | 0.02* |
| GM volume | 0.46±0.01 | 0.41±0.01 | <0.001* | 0.41±0.01 | 0.42±0.01 | <0.001* | <0.001* | <0.001* | 0.07 |
| WM volume | 0.35±0.01 | 0.34±0.01 | <0.001* | 0.34±0.01 | 0.35±0.01 | <0.001* | <0.001* | 0.01* | 1 |
| CSF volume | 0.18±0.01 | 0.23±0.01 | <0.001* | 0.24±0.01 | 0.23±0.01 | <0.001* | <0.001* | <0.001* | 0.02* |

## 3. Results

### 3.1. Age-related and education-related differences of socio-demographic variables and brain measures

The analysis of socio-demographic variables revealed that the young group differed from the old group in terms of BMI and sex (Table 1). VBM analysis showed widespread reduction in GM volume in the old group compared to the young group. Brain regions with group differences included the anterior cingulate, precuneus, superior, middle and inferior frontal gyri, supramarginal gyrus, insular cortex, temporal, parietal and lateral occipital regions bilaterally (Figure S1 in Supplementary Material). These results are in line with previously reported findings for this type of comparison (Filippini et al., 2012). No voxel-wise structural differences were observed between old-high and old-low participants. For completeness and descriptive purposes, we presented further structural MRI data analysis showing that the young group had higher TBV, GM, and WM, and lower CSF values, compared to the old group (Table 1). Differences were found between the old-high and the old-low groups in terms of TBV and CSF.

### 3.2. Age-related and education-related differences in cognitive performance and functional connectivity

The analysis of cognitive scores showed that the young group performed better than the old group in all the cognitive tests, apart from vocabulary (Table 2). For this variable, the old group did not differ from the young group. However, when splitting the older participants according to their educational level, the old-high, who had both higher word knowledge and higher education, showed the largest vocabulary size (p<0.01). Overall, the old-high group showed lower scores on cognitive tests as compared to the young group, but higher scores as compared to the old-low group, see Table 2. Moreover, the old-high participants did not differ from the young participants in two cognitive tests requiring the use of high-level cognitive function (i.e., verbal fluency), see Table 2. The rs-fMRI analysis revealed both age- and education-related differences in functional connectivity values. From the derived 25 components representing temporally correlated fMRI signals in different brain regions, we were able to identify eight spatial maps showing topological patterns associated with well-known RSNs (Figure 1). The RSNs included Somatomotor network (SMN), Visual-Medial network (VMN), Visual-Peripheral network (VPN), Salience network (SN), Dorsal Attention network (DAN), Default Mode network (DMN), right Fronto-Parietal network (rFPN), and left Fronto-Parietal network (lFPN).

**Table 2.** Descriptive analyses and Analysis of Covariance (ANCOVA) of Cognitive performance and Resting State Networks connectivity strength for all the groups, i.e., younger (Y), older (O), older with higher level of education (OH), and older with lower level of education (OL). The table shows for each variable mean and standard deviation. Columns show the values of the descriptive statistics for each group separately, i.e. young, old, old-high, and old-low, and for each comparison between groups, i.e. young vs old (third column), young vs old-high (seventh column), young vs old-low (eighth column), and old-high vs old-low (ninth column) contrasts. The Bonferroni-corrected p-values associated with the one-way ANCOVA (sixth column) are also reported for each contrast. The results of the ANCOVA and post-hoc comparisons are corrected for the effect of TBV BMI, and sex. Asterisks indicate significant results.

|  | Y<br>(N=137) | O<br>(N=60) | Y vs O<br>*p-val* | OH<br>(N=30) | OL<br>(N=30) | *p*-ANCOVA | Post-Hoc comparisons<br>(Bonferroni-corrected) |  |  |
| --- | --- | --- | --- | --- | --- | --- | --- | --- | --- |
|  |  |  |  |  |  |  | Y vs OH | Y vs OL | OH vs OL |
|  | Cognitive Performance |  |  |  |  |  |  |  |  |
| Immediate Memory Recall | 13.43±2.39 | 10.90±2.90 | <0.001* | 10.66±3.16 | 11.13±2.66 | <0.001* | <0.001* | <0.001* | 1 |
| Delayed Memory Recall | 13.14±2.43 | 10.13±2.67 | <0.001* | 9.77±2.94 | 10.50±2.35 | <0.001* | <0.001* | <0.001* | 1 |
| Phonemic Fluency | 24.70±6.40 | 21.28±5.23 | <0.001* | 21.97±5.89 | 20.60±4.46 | <0.01* | 0.10 | <0.01* | 0.85 |
| Semantic Fluency | 41.19±9.04 | 35.90±9.65 | <0.001* | 37.70±8.38 | 34.10±10.61 | <0.01* | 0.41 | <0.01* | 0.41 |
| Vocabulary | 33.26±2.68 | 33.63±2.57 | 0.37 | 34.83±2.40 | 32.43±2.17 | 0.001* | <0.01* | 1 | <0.01* |
|  | Resting State Networks - connectivity strength |  |  |  |  |  |  |  |  |
| SMN | 17.23±8.28 | 17.75±9.84 | 0.69 | 18.92±11.27 | 16.58±8.20 | 0.84 | 1 | 1 | 1 |
| VMN | 27.62±9.12 | 24.01±9.01 | 0.01* | 23.63±10.30 | 24.39±7.66 | 0.03* | 0.03* | 0.17 | 1 |
| VPN | 11.40±6.97 | 13.38±5.51 | 0.05 | 12.75±4.12 | 14.03±6.64 | 0.38 | 1 | 0.52 | 0.90 |
| SN | 14.63±4.75 | 16.65±5.73 | 0.009* | 17.05±7.01 | 16.26±4.14 | 0.09 | 0.23 | 0.11 | 1 |
| DAN | 12.73±2.90 | 10.62±3.25 | <0.001* | 11.32±3.44 | 9.92±2.94 | <0.001* | 0.04* | 0.001* | 0.77 |
| DMN | 18.03±3.56 | 16.16±4.28 | 0.002* | 15.75±5.06 | 16.56±3.37 | 0.04* | 0.02* | 0.52 | 0.88 |
| rFPN | 12.95±3.34 | 12.46±5.86 | 0.39 | 13.05±4.70 | 11.86±2.71 | 0.64 | 1 | 1 | 1 |
| IFPN | 12.45±3.29 | 11.26±4.51 | 0.04* | 11.42±5.30 | 11.11±6.63 | 0.36 | 0.42 | 1 | 1 |

**Figure 1.**
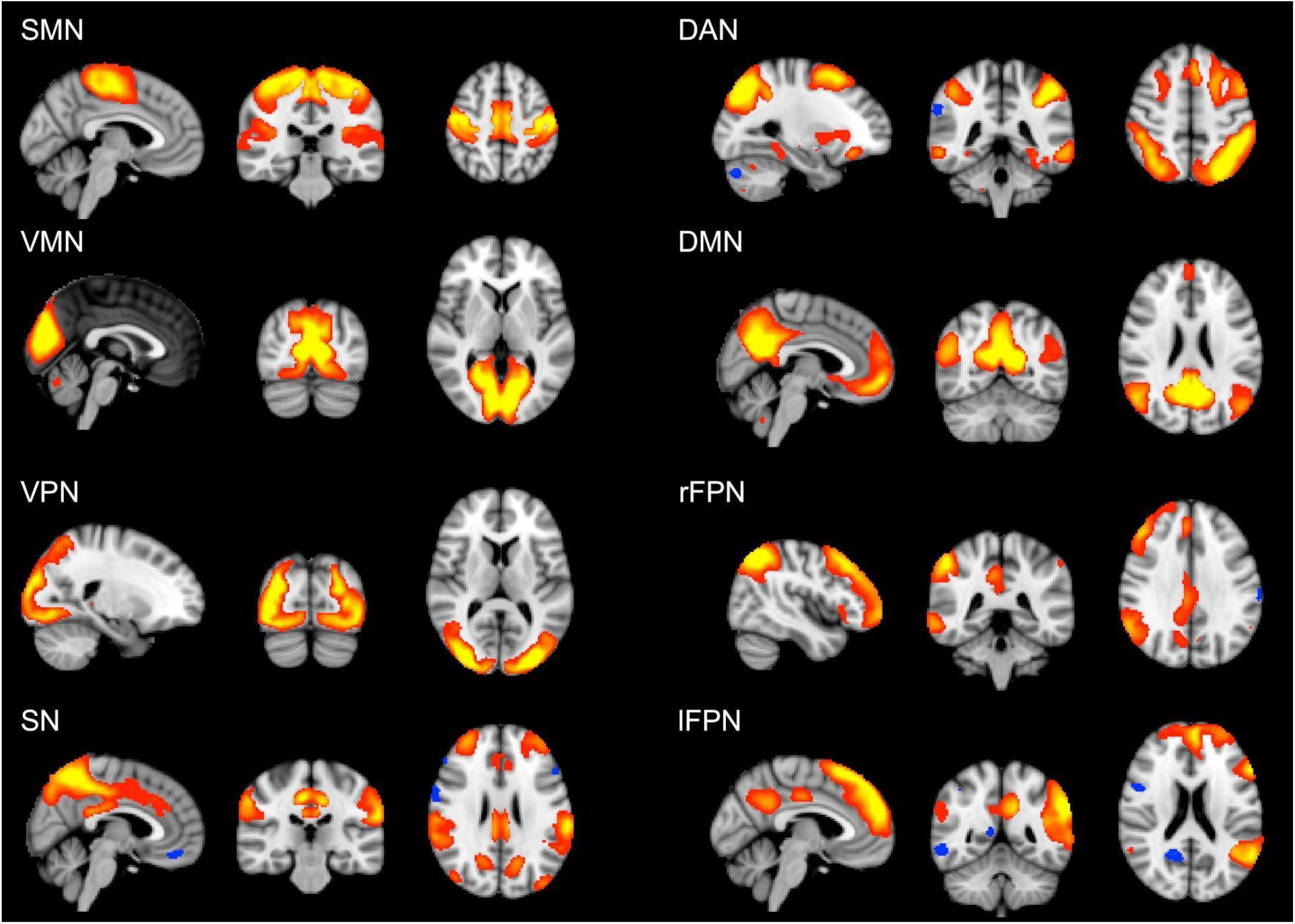
Large-scale brain networks reconstructed using rs-fMRI data considering all the subjects. RSNs were selected and labeled following visual check as: Somatomotor network (SMN), Visual-Medial network (VMN), Visual-Peripheral network (VPN), Salience network (SN), Dorsal Attention network (DAN), Default Mode network (DMN), right Fronto-Parietal network (rFPN), and left Fronto-Parietal network (lFPN). Group-level RSN maps (N = 197) are thresholded at z = 3. Red to yellow colors represent z-scores >3.

Overall, t-tests showed significant differences in connectivity strength, between the young and the old group for the VMN, the SN, the DAN, the DMN, and the lFPN (Table 2). By splitting the old group based on education, in ANCOVA, we identified some specific findings. In particular, the VMN showed that the old-high group had significantly lower network connectivity strength compared to the young group (*p* = 0.03), which was not reported in the case of the old-low group. Similarly, in the DMN, the old-high group showed significantly lower connectivity strength compared to the young group (*p* = 0.02), which was not reported in the case of the old-low group. Differently from the above reported RSNs showing significant differences between the young and the old group, the DAN showed significant differences for both the old-high and the old-low groups compared to the young group (old-high vs young *p* = 0.04; old-low vs young *p* = 0.001) also in the post-hoc comparisons.

### 3.3. Association between cognitive performance and resting state network connectivity strength: an exploratory correlation analysis

An exploratory correlation analysis between network connectivity strength and cognitive scores was performed for those RSNs revealing significant differences between groups, i.e., the VMN, the DAN, and the DMN (Table 2). The correlation analysis showed a significant negative relationship between the DMN connectivity strength and the memory scores (Table S1 in Supplementary Material). The Fisher’s-r-to-Z transformation of the Pearson’s correlation relative to the DMN-memory showed that the correlation slopes of the young and the old groups were significantly different between each other, in both Immediate Memory (two-tailed *p* = 0.02) and Delayed Memory recall (two-tailed *p* = 0.02). Such differences were substantially carried out by the old-high group, which showed preserved significance (two-tailed *p* = 0.04) for the Delayed Memory recall and approached significance (two-tailed *p* = 0.06) for the Immediate Memory. The relationship between both memory scores and the DMN connectivity strength were clearly not significant in the old-low group Indeed, the differences between correlation slopes approached a significant result in the comparison between the old-high and the young groups (two-tailed *p* = 0.05), while they were clearly non-significant in the comparison between the old-high and the old-low groups (two-tailed p = 0.20), see Figure 2.

**Figure 2.**
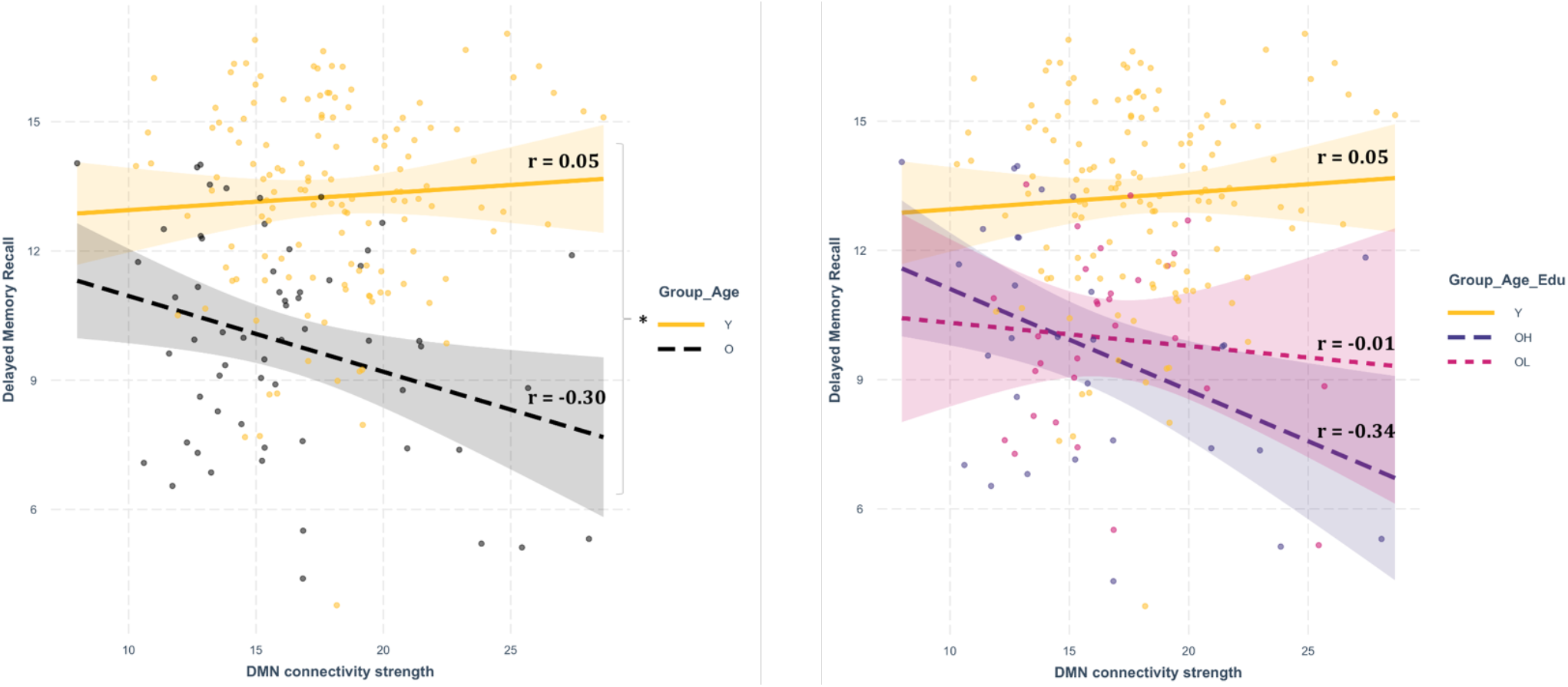
Correlation Analysis (Pearson’s Correlation) between DMN connectivity strength and Delayed Memory recall. The panel on the left-hand side shows the slopes, the residuals and the Pearson’s r associated with the partial correlation analyses between DMN connectivity strength and the Memory score, in the young and the old groups. The panel on the right-hand side shows the slopes, the residuals and the Pearson’s r associated with the partial correlation analyses between DMN connectivity strength and the Memory score in the young, the old-high, and the old-low groups. The asterisk indicates the result of significant two-tailed differences between slopes.

## 4. Discussion

In this study, we investigated whether education differentiates cognitive performance and resting state fMRI connectivity in healthy aging. To this end, we analyzed cognitive, structural and functional brain imaging variables and we considered education like a crucial socio-demographic aspect. Older adults typically show cognitive, structural, and functional age-related differences compared to younger adults (e.g., lower accuracy and slower response times in cognitive tasks, as also cortical thinning and brain atrophy) as the possible result of a reduced neural activity (see in Festini, 2018). However, this is not always the case, as brain overactivation has also been reported in healthy aging, in response to a compensatory necessity (e.g., Cabeza, 2002). Accordingly, age-related differences at the cognitive level are not always associated with the differences at brain structural and functional level. These differences might also be linked to education, which has been used in previous literature as a predictor of cognitive performance in healthy aging (Montemurro et al., 2021). Notably, education helps to estimate resources possibly accumulated from early life, that may support cognition also in later life (Mondini et al., 2022), although some studies have reported different findings (Nyberg et al., 2012). For this reason, we wanted to analyze whether cognitive and neural variables in aging could be further specified whereby a difference at the level of education is taken int account. In this study, the variable education did not vary within the young adults, who were all highly educated (i.e., *gymnasium*), but varied within the older adults, who were divided in two subgroups based on their level of education (i.e., *gymnasium* and *realschule* for old-high and old-low groups, respectively). With this procedure, we first examined whether there were any age-related differences between young adults and older adults. Then, we investigated whether education-specific features were identified in the old-high and the old-low groups, which is the core novelty of this study.

In line with previous literature, younger adults performed better than older adults in all cognitive tests, except for the vocabulary in which older adults performed better than younger adults (Bettio et al., 2017; Nyberg and Pudas, 2019). Importantly, the young and the old-high groups did not differ between each other in terms of language performance (i.e., verbal fluency tests) as a possible effect of education (Barulli and Stern, 2013) but the old-high group had a significantly higher vocabulary score compared to the other two groups. Overall, these results confirmed that higher education may provide for a more “youth-like” performance on language tasks requiring flexibility and cognitive control. In addition, vocabulary may have increased both along the lifespan and through the education level, which may have played as a sort of reserve/storage of verbal capacities too (see also the DACHA theoretical model in (Verhaeghen, 2003; Spreng and Turner, 2019).

We found between-group differences in terms of brain structure, with the older adults showing signs of deterioration, i.e., reduced gray and white matter, increased CSF, compared to young adults (Fjell et al., 2009; Salthouse, 2011; Orellana et al., 2016; Oschwald et al., 2019), regardless the level of their education. From a functional point of view, the derived RSNs (i.e., SMN, VMN, VPN, SN, DAN, DMN, rFPN and lFPN) presented a more heterogeneous scenario. A significantly higher RSN connectivity strength was found in the young group compared to the old group in the VMN, the DAN, and the DMN, which notably may be affected by aging (Onoda et al., 2012; Jockwitz et al., 2017; Farras-Permanyer et al., 2019), while the SN showed an opposite trend (higher connectivity strength for the older adults, especially for those with a high level of education). When accounting for education, the results significantly differed in the RSN connectivity strength for the VMN, the DAN, and the DMN (Table 2). In the DAN, the highest level of network connectivity strength was found for the young group, followed by the old-high, and then the by the old-low group. Interestingly, in the DAN, both the old-high and the old-low groups significantly differed from the young group, which suggested a clear age-related effect for this network (Jockwitz and Caspers, 2021). For both the VMN and the DMN, the old-high group had the lowest network connectivity strength in line with previous research that showed an age-related decreased connectivity of these RSNs (Farras-Permanyer et al., 2019; Jockwitz and Caspers, 2021) which in turn display the potential compensatory effects carried out by education. These findings are also in line with the STAC-r model (Reuter-Lorenz and Park, 2014), which highlighted education for explaining compensation processes in aging.

The results of this study suggest that higher education makes older individuals possibly more cognitively specialized in terms of verbal abilities, which is shown by the highest vocabulary skills in the old-high group compared to the other two groups, and by the non-significant differences between the old-high and the young group on verbal fluency (which also requires executive control). This suggests that a higher education defines specific neural patterns in the brain at rest. Critically, the old-high group not only displayed the lowest connectivity strength in the DMN, but, in this group, the DMN was the only network showing a selective, negative, association with memory capacities (Table 3). Although the correlation analyses were only observational and exploratory, we may explain these results by underlining that the DMN, which is known to “deactivate” proportionally with the increasing cognitive demand (Anticevic et al., 2012), may reflect a peculiar compensatory effect carried out by education at the level of memory capacities, which are typically vulnerable to aging (Farras-Permanyer et al., 2019). Age-related dysregulation in the connectivity of the DMN has been previously found in association with compensatory mechanisms, expressed in terms of recruitment of alternative brain areas (Park and Reuter-Lorenz, 2009; Reuter-Lorenz and Park, 2010). Moreover, our findings fit with the DECHA theoretical model (i.e., Default-Executive Coupling Hypothesis of Aging, (Spreng and Turner, 2019), which has more recently been developed for investigating complex compensatory mechanisms in aging. According to DECHA, education might help in the shift toward a more “semanticized” cognition (Spreng and Turner, 2019), which in turn would explain how the old-high group in this study have also better coped with cognitive demands in the semantic verbal fluency tasks compared with the old-low group.

The cross-sectional nature of this dataset represents a limitation in this study. Longitudinal studies would be warranted to properly study age-related differences by detecting changes in brain structure and function over time (Damoiseaux, 2017). Furthermore, criteria used to define the level of education might be considered to some extent suboptimal, as we could not use the continuous education values but two broad classes. However, such limitations were not specifically linked to the experimental choices of this study but represent intrinsic features of the database (LEMON).

## 5. Conclusion

Education maintains its modulatory role in neurocognitive functioning. It is able to differentiate cognitive and neural profiles of healthy older adults, based on their high-order verbal proficiency and resting-state connectivity of neural networks typically employed in high-order cognitive functioning. A key network in this context is the DMN as it might reflect memory compensatory mechanisms in older adults with higher education, who may have, at the cognitive level, a higher potential amount of resources to employ in their every-day lives.

## AUTHOR CONTRIBUTIONS

**Sonia Montemurro**: Conceptualization, Data Curation, Formal Analysis, Writing – original draft; **Nicola Filippini**: Conceptualization, Formal Analysis, Writing – review & editing; **Giulio Ferrazzi**: Data Curation, Formal Analysis, Writing – review & editing; **Dante Mantini**: Funding Acquisition, Writing – review & editing; **Giorgio Arcara**: Conceptualization, Funding Acquisition, Writing – review & editing; **Marco Marino**: Conceptualization, Data Curation, Formal Analysis, Funding Acquisition, Writing – original draft

## ACKNOWLEDGEMENTS

A special thanks to the Day Clinic for Cognitive Neurology of the University Clinic Leipzig and the Max Planck Institute for Human and Cognitive and Brain Sciences (MPI CBS) in Leipzig, Germany. Funding: MM is funded by Research Foundation Flanders (FWO) (postdoctoral fellowship 1211820N)

